# Prediction errors for aversive events shape long-term memory formation through a distinct neural mechanism

**DOI:** 10.1101/2021.03.19.436177

**Authors:** Felix Kalbe, Lars Schwabe

**Author notes:** Corresponding author: Lars Schwabe, Institute of Psychology, Universität Hamburg, Von-Melle-Park 5, 20254 Hamburg, Germany.

## Abstract

Prediction errors (PEs) have been known for decades to guide associative learning, but their role in episodic memory formation has been discovered only recently. Using an encoding task in which participants learned to predict which stimuli are followed by aversive shocks, combined with univariate, multivoxel, and large-scale network analyses of fMRI data, we show that enhanced memory for events associated with negative PEs was linked to reduced hippocampal responses to PEs and increased crosstalk between the ‘salience network’ and a frontoparietal network commonly implicated in memory formation for events that are in line with prior expectation. These PE-related effects could not be explained by mere changes in physiological arousal or the prediction itself. Our results suggest that superior memory for events associated with high PEs is driven by a distinct neural mechanism that might serve to set memories of high PE events apart from those with expected outcomes.

## INTRODUCTION

Imagine meeting Barack Obama in the supermarket. Most likely, this event would deviate strongly from what you expected during your grocery shopping. Such a mismatch between expectation and experience is referred to as a prediction error (PE). PEs play a key role in many cognitive domains, from perception to attention or language (den Ouden et al., 2012; Rao & Ballard, 1999; Spratling, 2008), and the comparison of predictions against perceptual evidence has been proposed as a unifying principle of neural computation (Bar, 2009; Friston, 2010). Traditionally, PEs are considered a to be driving force in reinforcement learning, during which an organism learns incrementally how to achieve pleasant and avoid unpleasant states (Glimcher, 2011; Niv, 2009). Beyond driving incremental learning across multiple instances, it may be expected that single episodes encoded in the context of a high PE, such as meeting Barack Obama by chance, are also preferentially stored in episodic memory. Although the preferential storage of high-PE events in long-term memory would aid behavioral adaptation and optimal choice in the future (Gershman & Daw, 2017; Shohamy & Adcock, 2010), PEs received surprisingly little attention in episodic memory research (for early exceptions, see Henson & Gagnepain, 2010; Mizumori, 2013). Only very recently, behavioral evidence started to accumulate showing that PEs may extend into episodic memory and promote memory formation of nearby events (Ergo et al., 2020; Greve et al., 2017; Jang et al., 2019; Kalbe & Schwabe, 2020; Rouhani et al., 2018). A fundamental question concerns how PEs boost long-term memory formation.

One way through which PEs may promote episodic memory for surrounding events is by enhancing well-known mechanisms of long-term memory formation. Since the seminal case of H.M. (Scoville & Milner, 1957), memory formation has been strongly linked to the medial temporal lobe, including the hippocampus and adjacent cortices (Alvarez & Squire, 1994; Eichenbaum, 2001; Moscovitch et al., 2016). It is further well established that hippocampal memory formation is enhanced by emotional arousal, which is likely to be the result of a high PE event such as meeting Barack Obama (de Berker et al., 2016). This emotional memory enhancement is thought to be mediated by the amygdala, which strengthens memory formation processes in the hippocampus and related areas that together form a “medial temporal encoding network” (MTEN) (Hermans et al., 2014; McGaugh & Roozendaal, 2002; Richardson et al., 2004; Strange & Dolan, 2004). Thus, one hypothesis would be that PE-driven episodic memory enhancements are due to increases in medial-temporal lobe activation.

Alternatively, PEs might drive long-term memory formation through mechanisms that are critically distinct from those known to underlie common memory formation. Initial behavioral evidence suggests that PE effects on episodic memory formation go beyond the effects of physiological arousal (Kalbe & Schwabe, 2020), suggesting that PE effects may not be fully attributable to mere quantitative enhancements of standard memory processing.

Furthermore, events associated with high PEs have been suggested to create event boundaries in memory and therefore disrupt the structure of memories by establishing a new latent context resulting in a separate memory trace (Rouhani et al., 2020). These behavioral findings point to the alternative that PEs might induce a qualitative shift in mnemonic processing, towards more exemplar-based representations and by setting the PE-associated events apart from those with expected outcomes. When an event is associated with a large PE, this triggers an alertness response to allocate attentional resources to the unexpected outcome (Metereau & Dreher, 2013; Summerfield & Egner, 2009). This neural response may be mediated by the salience network (Fouragnan et al., 2018; Ham et al., 2013), which is known to play a key role in attentional capture of biologically relevant events and is mainly comprised of the bilateral anterior insula and the dorsal anterior cingulate cortex (dACC; Garrison et al., 2013; Ham et al., 2013). At the same time, if high PE events are processed separately from existing knowledge structures represented in what is referred to as a schema (Ghosh & Gilboa, 2014), it can be further predicted that PEs result in a decreased recruitment of the neural ‘schema-network’, comprised mainly of the angular gyrus, the precuneus, and the medial prefrontal cortex (mPFC; van Kesteren et al., 2012; Vogel et al., 2018a). Accordingly, this alternative view predicts that the enhanced memory for events encoded in the context of high PEs is due to an activation of the salience network, accompanied by an even reduced activation of areas implicated in memory formation for events that are in line with prior experience (i.e., the MTEN and ‘schema-network’).

To test these alternative hypotheses and provide insight into the mechanism through which PEs can enhance episodic memory, we had participants perform an incidental encoding task in which they saw a series of stimuli from different categories that were associated with different probabilities to receive a mild electric shock. Participants were asked to predict the occurrence of a shock on a trial-by-trial basis. Comparing these shock expectancy ratings to the actual trial outcome indicated whether participants experienced a negative PE (no shock occurred when it was expected to occur), a positive PE (a shock occurred that was not predicted), or no PE at all (Delgado et al., 2008; McHugh et al., 2014; Miller et al., 1995; Schultz, 1998). Memory was probed in a recognition test 24 hours after encoding. To unravel the neural mechanisms underlying PE-related enhancements of episodic memory, we used behavioral modelling, autonomic arousal measurements, and fMRI in combination with multivoxel pattern analysis (MVPA) and large-scale network analysis.

## RESULTS

To test how PEs shape long-term memory formation, 50 participants completed a combined incidental encoding and fear learning task in an MRI scanner (Figure 1A). In this task, consisting of four runs with 30 trials each, participants saw a random sequence of unique pictures from three distinct categories (i.e., vehicles, clothing, and tools) and received an unpleasant electric shock after some pictures but not others. On each trial, participants were required to estimate the probability that a shock would follow on a scale from 0 to 100 percent. Critically, each picture category was linked with a pre-defined shock probability of either 67% (CS^a+^), 33% (CS^b+^), or 0% (CS^-^), which resulted in a different magnitude (and direction) of PEs associated with these stimulus categories. Explicit shock expectancy ratings also allowed us to evaluate participants’ uncertainty in each trial, expressed as the closeness of the rating to 50%. In addition to shock expectancy ratings, we recorded anticipatory (i.e., CS-induced) and outcome-related skin conductance responses (SCR) as a measure of physiological arousal. About 24 hours after the encoding session, participants completed a surprise recognition test that included all pictures shown during encoding and the same number of new pictures from each of the three categories.

**Figure 1.**
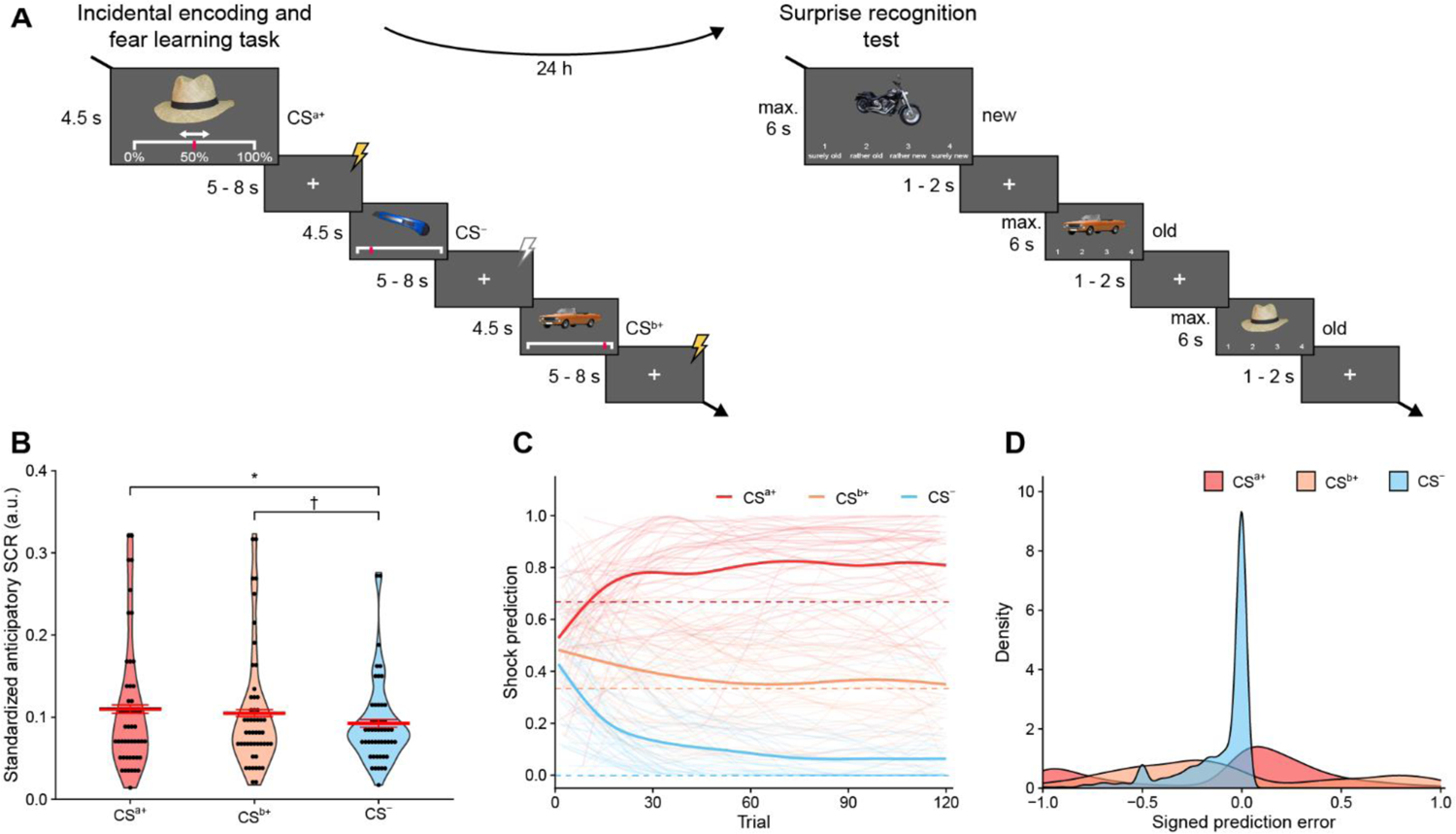
Experimental task and performance parameters (A) Participants completed a combined incidental encoding and fear learning task and a surprise recognition test for its contents about 24h later. In the encoding task, participants saw a series of unique pictures from three different categories (clothing, vehicles, and tools) linked to fixed probabilities to receive an electric shock (CS^a+^ − 67%, CS^b+^ − 33%, and CS^-^ − 0%). On each trial, participants indicated their shock expectation. Approximately 24 h later, they saw all pictures from the previous day intermixed with the same number of new pictures and categorized each picture as either ‘old’ or ‘new’. (B) Mean standardized anticipatory skin conductance responses (SCR) confirmed successful fear conditioning, as reflected in significantly elevated SCR to both CS^a+^ and CS^b+^ items compared with CS^-^ items. Black dots show data from individual participants. Thick red bar represents group mean, while thin red bars show ± 1 standard error of the mean. (C) Participants’ mean shock expectancy ratings (thick lines) approached the true shock probabilities (dotted lines) relatively fast, although there was a tendency to overestimate shock probabilities. Thin lines represent data from individual participants. (D) Signed PEs were distributed relatively symmetrical for CS^a+^ and CS^b+^ pictures around zero. PEs for CS^-^ pictures were mostly zero, reflecting that participants learned that items from this category were never paired with a shock. † p<.05, * p_corr_<.05 (Bonferroni-corrected).

### Successful fear learning

Physiological and explicit rating data indicated successful fear learning. Specifically, standardized anticipatory SCR differed significantly between CS categories, F(2,98)=3.62, p=.030, ηG=.011 (Figure 1B). Post-hoc paired t-tests revealed that participants showed increased anticipatory SCRs to both CS^a+^ pictures (t(49)=2.38, p_corr_=.042 (Bonferroni-corrected), d_av_=0.27) and CS^b+^ pictures compared with CS^-^ pictures (t(49)=2.10, p=.041, p_corr_=.082 (Bonferroni-corrected), d_av_=0.20). Explicit shock ratings further showed that participants learned to associate picture categories with their respective shock probabilities over the course of the task (Figure 1C). Participants had a significantly higher shock expectancy for CS^a+^ than for CS^b+^ (t(49)=11.53, p_corr_<.001 (Bonferroni-corrected), d_av_=2.67) and for CS^b+^ than for CS^-^ (t(49)=10.81, p_corr_<.001 (Bonferroni-corrected), d_av_=1.87).

From participants’ explicit shock expectancy ratings, we derived signed PEs by contrasting each prediction with the binary outcome (i.e., unshocked or shocked) in the respective trial (see Methods). Resulting PEs ranged from −1 to 1, with negative values in cases of unexpected shock omissions and positive values in cases of unexpected shocks, while greater distances from 0 in both directions indicated greater discrepancies between predictions and outcomes. Importantly, the distribution of signed PEs varied almost symmetrically around 0 (Figure 1D), allowing similarly reliable conclusions about effects of both negative PEs and positive PEs. Moreover, the explicit shock ratings allowed us to directly assess participants’ prediction uncertainty, which ranged from 0 (maximal certainty, corresponding to predictions of 0% or 100%) to 1 (maximal uncertainty, corresponding to a prediction of 50%).

### Overall recognition memory performance

In the recognition test 24 hours after encoding, participants performed overall very well, as indicated by markedly higher hit rates (i.e., the rate of correctly classifying previously seen pictures as ‘old’) than false alarm rates (i.e., the rate of incorrectly classifying unseen pictures as ‘old’), Mhitrate=60.9% (SD=0.149), MFArate=21.1% (SD=0.098). Participants were significantly more certain with their responses for hits (M=0.59; SD=0.18) than for false alarms (M=0.26, SD=0.20), t(49)=15.92, p<.001, d_av_=1.70.

A repeated-measures ANOVA showed that hit rates differed significantly between CS categories, F(2,98)=7.29, p=.001, ηG²=0.05. For false alarm rates, on the other hand, there was no such difference between CS categories, F(2,98)=0.25, p=.77, ηG²=.003, suggesting that the actual memory but not the response bias differed between CS categories. Post-hoc paired t-tests showed that hit rates were selectively enhanced for items from the CS^a+^ category, which was associated with a shock probability of 67%, compared with both items from the CS^b+^ category (t(49)=4.15, p_corr_<.001 (Bonferroni-corrected), d_av_=0.54), which was associated with a shock probability of 33%, and the CS^-^ category (t(49)=2.64, p_corr_=.022 (Bonferroni-corrected), d_av_=0.40; Figure 2A), which was never followed by a shock. Enhanced recognition performance for CS^a+^ items was also obtained when hits and false alarms were integrated to the sensitivity d’ based on signal detection theory: A repeated measures ANOVA confirmed that d’ was generally different between CS categories (F(2,98)=3.70, p=.028, ηG²=.03), with post-hoc t-tests confirming an increased memory sensitivity for CS^a+^ items compared with both CS^b+^ items (t(49)=2.28, p=.027, p_corr_=.053 (Bonferroni-corrected), d_av_=0.38) and CS^-^ items (t(49)=2.42, p_corr_=.038 (Bonferroni-corrected), d_av_=0.35).

**Figure 2.**
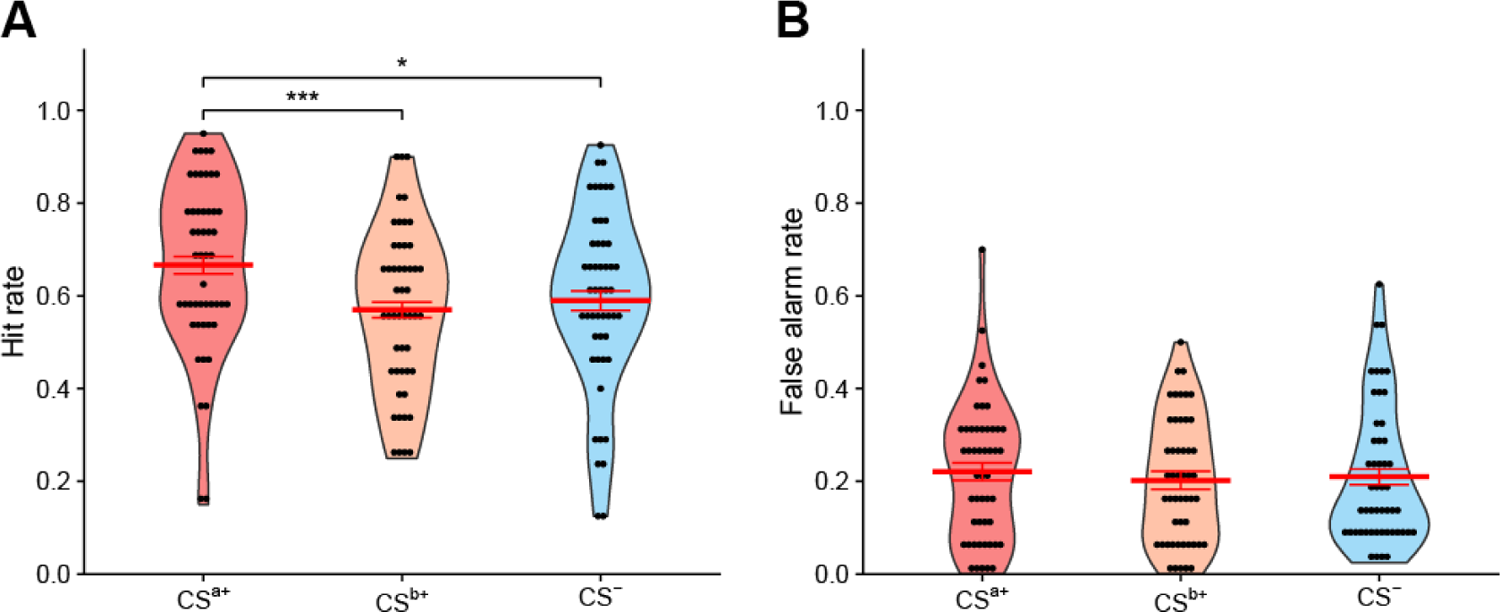
General recognition performance by CS category. (A) Hit rates for items from the CS^a+^ category were significantly larger compared with both CS^b+^ and CS^-^ items. Although CS^a+^ items had the highest shock probability, this could not explain their increased hit probability as hit rates for shocked items even tended to be lower than for unshocked items (supplementary Figure S1). (B) False alarm rates were comparable for all three conditioning categories, showing that the CS-type did not affect the mnemonic response bias. Black dots show data from individual participants. Thick red bar represent group means, while thin red bars show ± 1 standard error of the mean. * p_corr_<.05 (Bonferroni-corrected), *** p_corr_<.001 (Bonferroni-corrected)

At first glance, one might assume that these differences are simply due to differences in (arousing) shock presentations between CS categories. However, our data did not support this interpretation. The greater proportion of shocked items could not explain the improved hit rate for the CS^a+^ category: A repeated-measures ANOVA to explain hit rates indicated no memory advantage for shocked over unshocked items per se (F(1,49)=1.12, p=.294, ηG²=.022). Further, a 2×2 repeated-measures ANOVA confirmed increased hit rates for CS^a+^ over CS^b+^ items even after controlling for shocks (F(1,49)=19.47, p<.001, ηG²=.08). Notably, this ANOVA even showed a tendency towards decreased hit rates for shocked items (F(1,49)=3.76, p=.058, ηG²=.006), with no significant interaction (F(1,49)<0.001, p=.997, ηG²<.0001). These findings indicate that differences between CS categories in the number of presented shocks cannot explain the differential memory performance and that other factors drive the boost in memory.

### Aversive PEs and prediction uncertainty modulate episodic memory formation beyond arousal

To explain episodic memory formation in the incidental encoding task at trial level, we fitted generalized linear mixed-effects models (GLMMs) with a binary response variable (hit vs. miss) and a logit link function (i.e., mixed-effects logistic regression) using the *lme4 R* package (Bates et al., 2015). The dependent variable was the recognition of an item in the surprise recognition test, coded 0 for misses and 1 for hits. We applied the maximum random effects structure (Barr et al., 2013), estimating random intercepts and random slopes of all predictors per subject.

We fitted three initial models over all trials (including both negative and positive PEs) using (1) linear PEs, (2) quadratic PEs, and (3) a variant of quadratic PEs assuming that effects of negative vs. positive PEs would be in opposite directions based on the following inverted S-shaped transformation:

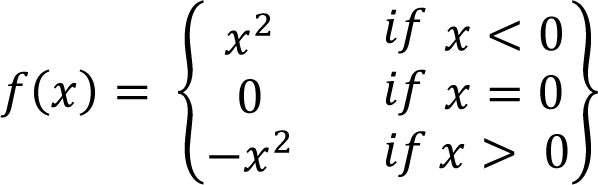

Model comparisons using the Akaike information criterion (AIC) to identify the optimal model while also considering increased model complexity favored the inverted S-shaped model (AIC = 7420.8) over both the linear (AIC=7425.2) and the quadratic model (AIC=7434.3). Results from this inverted S-shaped model indicated that negative PEs promoted memory formation, while positive PEs decreased memory formation, β=0.27, 95%-CI [0.07, 0.47], z=2.68, p=.007. Even after adding the binary occurrences of shocks to the model, this effect remained significant (β=0.58, 95%-CI [0.29, 0.87], z=3.30, p<.001), rejecting the notion that aversive shocks alone drive this effect.

Results so far suggest that greater negative PEs and greater positive PEs had opposite effects on episodic memory formation, with the former increasing and the latter decreasing the probability of a subsequent hit. However, this model assumes both effects to be equally strong in each participant. To further investigate whether this assumption is justified, we next fitted models separately for negative and positive PE trials with quadratic PEs as the sole independent variable to explain the binary recognition of an item (Figure 3A). For negative PEs, we again observed a memory enhancement with greater PE magnitude, β=0.49, 95%-CI [0.15, 0.82], z=2.85, p=.004. The same model for positive PEs confirmed that greater PE magnitude was instead associated with decreased memory performance, β=-0.73, 95%-CI [-1.12, −0.34], z=3.67, p<.001. Random βs per subject from both models were moderately negatively correlated, indicating that participants that showed a stronger memory benefit from negative PEs also showed a stronger memory decrease from positive PEs, r=-.395, t(48)=2.98, p=.005 (Figure 3B).

**Figure 3.**
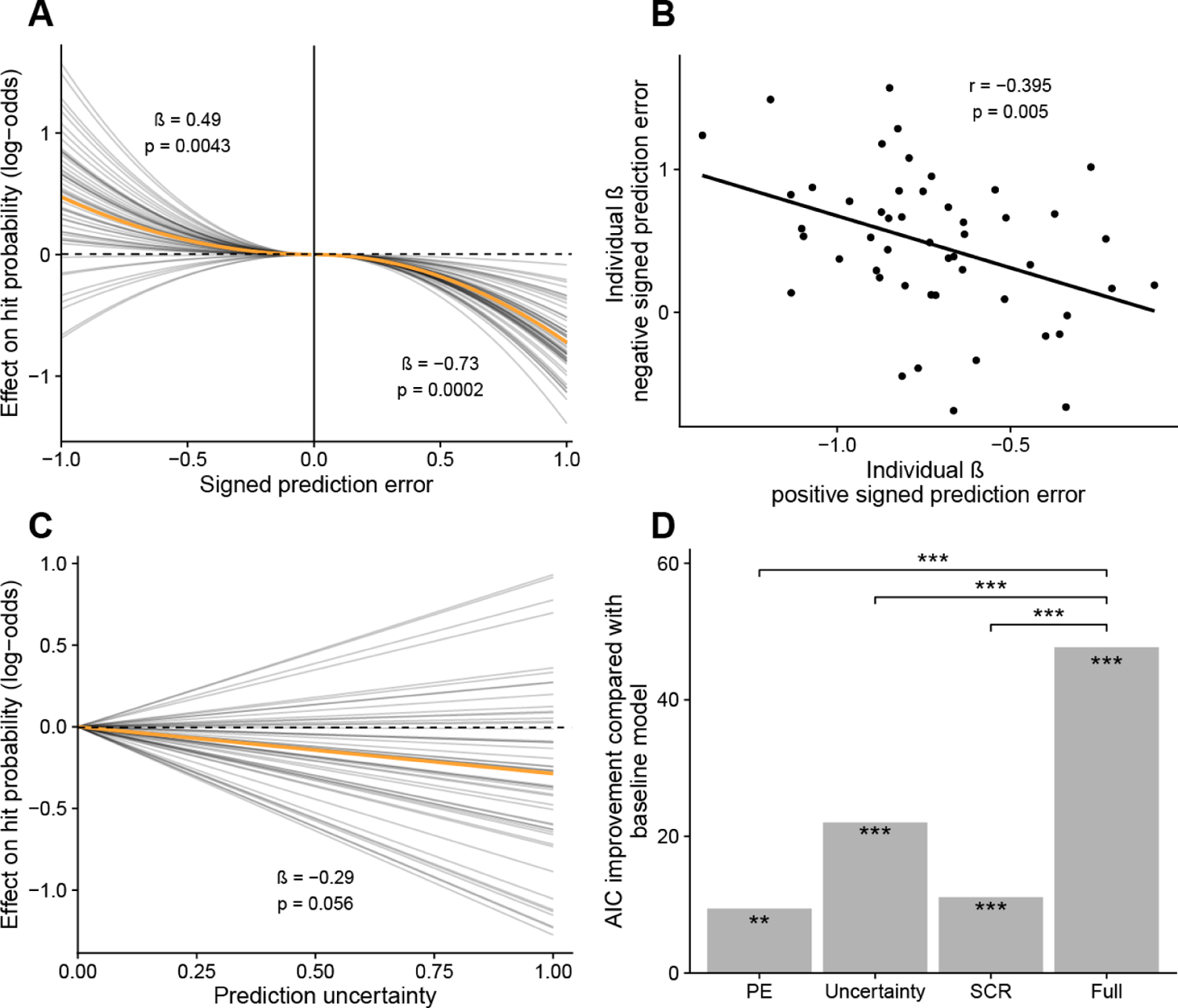
Behavioral model of long-term memory formation reveals modulating influences of prediction errors and prediction uncertainty (A) Results from a trial-level mixed-effect logistic regression show opposite effects of positive and negative prediction errors on later memory. Quadratic negative prediction errors (associated with unexpected shock omissions; left half) were linked with improved memory formation for associated pictures. In contrast, quadratic positive prediction errors (associated with shocks; right half) were linked with decreased memory formation. Orange line indicates estimated fixed effects of PEs, while thin black lines show PE effects estimated separately per participant. (B) Effects of quadratic negative and positive prediction errors were negatively correlated at the level of participants. (C) Prediction uncertainty was, independently of outcomes, associated with decreased hit probabilities. As in (A), the orange line indicates the estimated fixed effect, while thin black lines show participant-specific effect estimates. (D) Model comparisons showed that a full model combining PEs, uncertainty, and anticipatory and outcome-related SCRs explained memory formation better than any model containing only one of these measures. This was confirmed by both likelihood-ratio tests as well as the lowest (i.e., best) AIC value for this model. Notably, any model containing only a single predictor (i.e., either PEs, uncertainty, or SCRs) also performed significantly better than our baseline model comprised only of any a random intercept per participant (comparisons indicated by markings within each bar). ** p<.01, *** p<.001.

An alternative explanation for the memory-modulating effects of prediction errors could be that the mere expectation of an aversive shock drives the effect on long-term memory. If this were the case, then contrasting the explicit shock predictions with observed outcomes should not explain memory formation beyond the influence of mere predictions. To test this alternative account, we first fitted a mixed-effects logistic regression model with participants’ explicit shock predictions (ranging from 0 to 1) to explain the binary recognition of an item. Indeed, greater expectancy of a shock was associated with an increased probability that the item would subsequently be recognized, β=0.40, 95%-CI [0.12, 0.67], z=2.83, p=.005. After adding the subsequent binary occurrence of shocks, this positive effect of predictions remained (β=0.44, 95%-CI [0.16, 0.73], z=3.08, p=.002), while shocks themselves were not found to affect the probability that an item would be recognized (β=-0.04, 95%-CI [-0.20, 0.13], z=0.46, p=.65). To test whether PEs could explain memory formation beyond main effects of predictions and outcomes, we added them to the model using the inverted S-shaped transformation reported above. In contrast to the previous models, this model no longer indicated any memory-modulating effects of mere predictions, β=-0.67, 95%-CI [-1.53, 0.20], z=1.50, p=.13. However, aversive shocks were now associated with increased hit probabilities, β=0.83, 95%-CI [0.18, 1.47], z=2.51, p=.012. Most importantly, PEs again explained memory formation depending on their sign, β=1.22, 95%-CI [0.37, 2.08], z=2.81, p=.004. Further evidence that the model adding both outcomes and prediction errors fitted the data better compared with the prediction-only model came from a likelihood-ratio test, χ²(9)=26.72, p=.002. This was also reflected in a smaller AIC value for the model adding both outcomes and PEs (AIC=7366.7) compared with the prediction-only model (AIC=7375.4). Finally, as reported above, we did not find any improved recognition performance for items from the CS^b+^ category over items from the CS^-^ category, even though these were associated with significantly higher shock expectancies.

Closely related to both predictions and PEs, uncertainty about possible outcomes has been proposed to affect episodic memory formation as well (Stanek et al., 2019). In contrast to shock predictions, uncertainty is maximal when participants believe that the probability of a shock is 50% (i.e., maximal entropy). As before, we investigated effects of prediction uncertainty on memory formation using two mixed-effects logistic regression models with (1) linear uncertainty and (2) quadratic uncertainty as the sole independent variable to explain the binary recognition of an item. Because prediction uncertainty is independent of outcomes, these models included both unshocked and shocked trials. Results favored a linear model (AIC = 7408.2) over the quadratic model (AIC = 7418.2) and suggested a tendency towards a negative relationship between prediction uncertainty and memory formation for the associated item, β=-0.29, 95%-CI [-0.58, 0.008], z=1.91, p=.056 (Figure 3C). In other words, when participants were more uncertain in terms of their shock prediction, it tended to be less likely that they would later recognize the associated item.

Classic models of episodic memory formation in aversive contexts emphasized the memory promoting role of physiological arousal (Cahill & McGaugh, 1998; McGaugh, 2018). Therefore, we tested in a next step whether the PE-related memory changes that we observed here could be explained by physiological arousal. To this aim, we fitted a mixed-effects logistic regression model with standardized anticipatory SCRs and standardized outcome-related SCRs as the only two predictors for the binary recognition of an item. In this model, neither anticipatory SCRs (β=-0.14, 95%-CI [-0.58, 0.30], z=0.63, p=.531), nor outcome-related SCRs (β=0.26, 95%-CI [-0.11, 0.64], z=1.37, p=.170), had any significant effect on memory, suggesting that physiological arousal (expressed through SCR) did not drive long-term memory formation.

In a final mixed-effects logistic regression model, we included quadratic PEs, quadratic prediction uncertainty, anticipatory and outcome-related SCR in parallel to investigate whether previous results from simpler models would still hold after accounting for other memory-modulating variables. Since this model was again applied to trials including both negative and positive PEs, we again entered PEs using the previously introduced inverted S-shaped transformation. Results confirmed our previous findings that PEs had memory-promoting effects in the case of unexpected shock omissions and memory-decreasing effects in the case of unexpected shocks, β=0.44, 95%-CI [0.20, 0.67], z=3.62, p<.001. Further, this combined model confirmed our previous findings of memory decreasing effects of prediction uncertainty, β=-0.39, 95%-CI [-0.68, −0.09], z=2.56, p=.010. As before, standardized anticipatory SCRs had no significant effect on episodic memory formation, β=-0.17, 95%-CI [-0.62, 0.29], z=0.72, p=.47. Notably, unlike in the simpler SCR model, outcome-related SCRs showed a positive effect on memory formation, β=0.63, 95%-CI [0.18, 1.08], z=2.76, p=.006. Therefore, only after accounting for effects of PEs and uncertainty on memory formation, additional arousal-related influences occurred.

Separate model comparisons using likelihood ratio tests confirmed that the full model including PEs, uncertainty, and physiological arousal (measured by both anticipatory and outcome-related SCRs) was the most appropriate. This suggests that all three components uniquely and additively contribute to long-term memory formation. Compared with a simple baseline model containing only a random intercept for each participant, adding any type of predictor from the full model (i.e., either PEs, uncertainty, or physiological arousal) significantly improved the fit (all ps<.002; Figure 3D). Critically, the full model containing all three types of predictions led to the lowest AIC value. The full model also significantly improved the fit compared with any model containing only a single type of predictor (all ps<.001; Figure 3D).

### Negative PEs are associated with greater activation of the salience-network, paralleled by decreased activation of hippocampal and schema-network

Our behavioral findings show that negative PEs promote long-term memory formation, whereas positive PEs and uncertainty impair memory formation. To unravel the neural underpinnings of PE-driven changes in memory formation, we modelled the pre-processed fMRI time series using generalized-linear models (GLMs; see Methods). We aimed to identify neural correlates of previously identified memory modulators by specifying onsets of stimuli as a first regressor, with shock expectancy, and uncertainty as parametric modulators based on results from our behavioral models of memory formation. To control for the fact that these variables might also partially reflect arousal, we additionally added anticipatory SCRs as a third parametric modulator. A second regressor consisted of outcome onsets, with PEs as a parametric modulator based on results from our behavioral model. Again, we controlled for the possibly confounding effects of arousal with PEs by including outcome-related SCRs.

As for the PE-based behavioral models, we fitted this model separately for unshocked and shocked trials to account for the opposite effects of negative and positive PEs on memory formation that we observed. All of the findings reported below were significant at the whole-brain level at p<.05, FWE corrected. Negative PEs were associated with large clusters of increased activity in the bilateral anterior insula and the dACC, which are key regions of the salience network (Figure 4A-B) (Menon, 2011). In addition, negative PEs were associated with significant decreases in activation in large portions of the bilateral hippocampus (Figure 4D). Although it is important to note that the decrease in hippocampal activity occurred only after outcomes were revealed and therefore after the offset of the to-be-remembered stimulus, this finding is in stark contrast to earlier studies linking hippocampal activity to increased memory (Fernández et al., 1999; Shrager et al., 2008) and suggests that for the PE-induced memory enhancement that we observed here, a different mechanism may be at work. In addition to decreased activation in the hippocampus, we also observed decreased activity for negative PEs in the mPFC, precuneus, and left angular gyrus (Figure 4C-E), all three of which have been described as part of the schema network that links current information to existing knowledge structures (van Kesteren et al., 2012; Vogel et al., 2018a). This finding may be taken as first evidence that the superior memory for items associated with large negative PEs is associated with a distinct neural mechanism that sets these PE events apart from those with expected outcomes.

**Figure 4.**
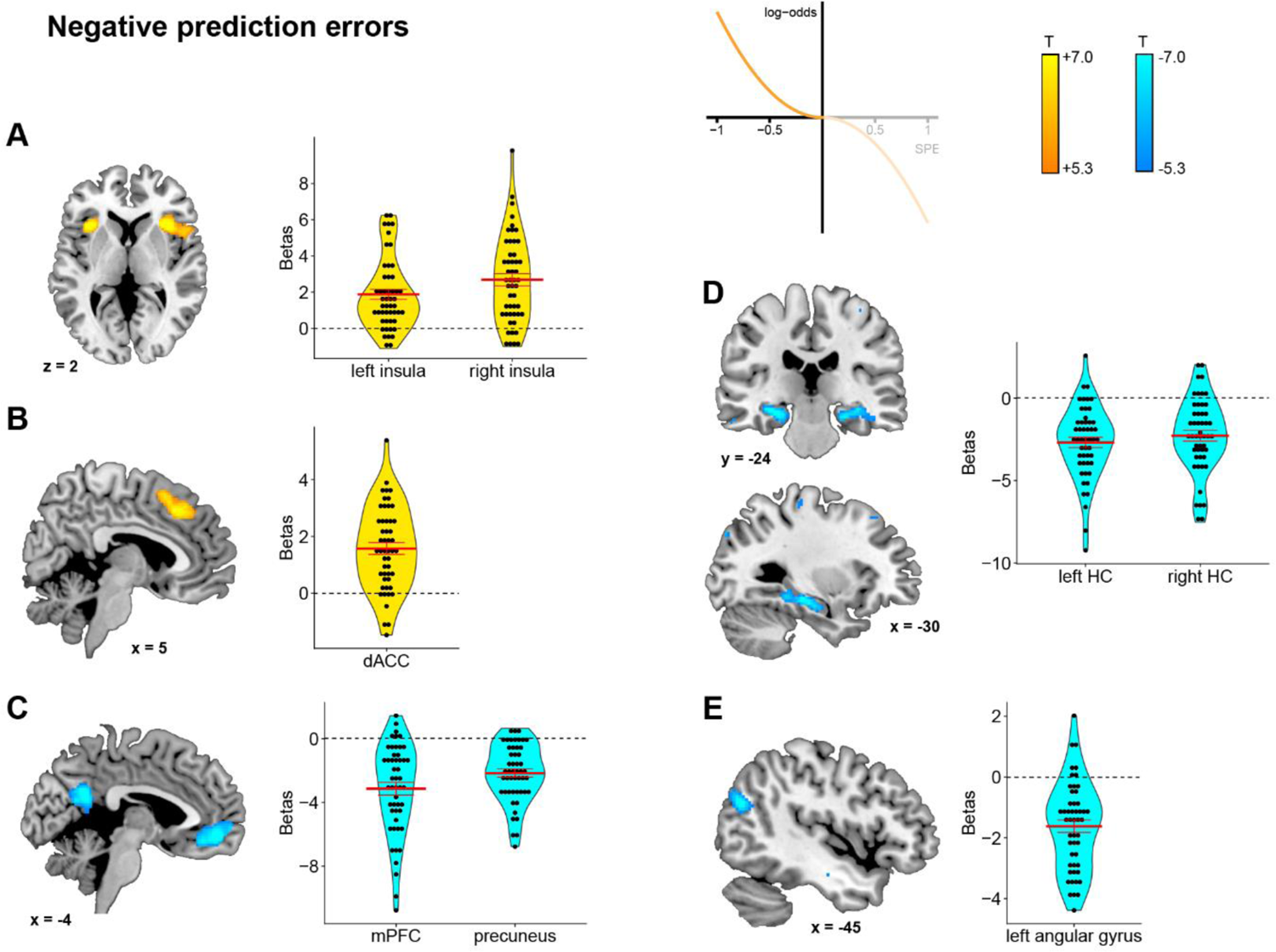
Univariate fMRI analysis to identify regions associated with negative PEs. Negative PEs were linked with increased BOLD responses in the bilateral insula and the dorsal anterior cingulate cortex (dACC), (A-B) and decreased BOLD-responses in the medial prefrontal cortex (mPFC), precuneus, bilateral hippocampus, and left angular gyrus (C-E). Only voxels significant at p < .05 after whole-brain family-wise error (FWE) correction (peak level) are displayed. Black dots indicate beta estimates from individual participants, while the red line shows the mean beta estimate over all participants.

Same as negative PEs, prediction uncertainty in unshocked trials was associated with decreased activation in the prefrontal cortex, although this cluster was located significantly more dorsally for uncertainty (supplementary Figure S2A). Additionally, we observed decreased activation in the bilateral middle temporal gyrus (supplementary Figure S2B), likely reflecting decreased visual processing of stimuli associated with greater prediction uncertainty, which might explain the reduced memory for items associated with uncertainty.

For mere shock expectancy, we found no significant changes in activation in any areas that were previously linked with PEs (i.e., dACC, insula, hippocampus, mPFC, precuneus, angular gyrus). Instead, shock expectancy was only associated with changes in occipital areas, which might reflect visual processing of the slider that participants used to give their expectancy rating (Supplementary Table 3). This finding complements results from the behavioral models suggesting that the deviations of outcomes from predictions (i.e., PEs) is critical for memory modulation, rather than the mere expectation of an aversive stimulus.

In parallel models for shocked trials, positive PEs were associated with increased activity in two smaller clusters located in the left superior parietal lobule and the right middle temporal gyrus and decreased activity in the left supramarginal gyrus. Prediction uncertainty in shocked trials was only associated with two small clusters of decreased activation in the right lingual gyrus and the left postcentral gyrus.

### Linking PEs with subsequent recognition

In a next step, we asked whether changes in brain activity in response to negative PEs would be predictive of the subsequent recognition of an item in the memory test. We specified a univariate fMRI model with onsets of unshocked outcomes as a regressor and PEs, the binary subsequent recognition of an item and their interaction as parametric modulators (see Methods). Based on the above findings and the existing literature, we identified three candidate regions of interest that may not only be associated with the signaling of aversive PEs but also linked to declarative memory formation (Bermudez-Rattoni, 2014; Eichenbaum, 2001; Euston et al., 2012): the bilateral insula, the ventromedial prefrontal cortex (vmPFC), and the bilateral hippocampus. Our analysis focused on the interaction between PEs and subsequent recognition, as this specific interaction links the processing of PEs with their effects on memory formation. Results showed that when larger negative PEs occurred and the item was subsequently recognized, we found decreased BOLD activity in the bilateral hippocampus, t(49)=1.41, p=.017, p_corr_=.052 (Bonferroni-corrected), d=0.35 (Figure 5). This finding provides further evidence that the memory-promoting effects of negative PEs are not simply due to an increase in ‘standard’ memory processing in medial-temporal regions. For the bilateral insula and the vmPFC there was no interaction between PEs and subsequent recognition (both ps>.16).

**Figure 5.**
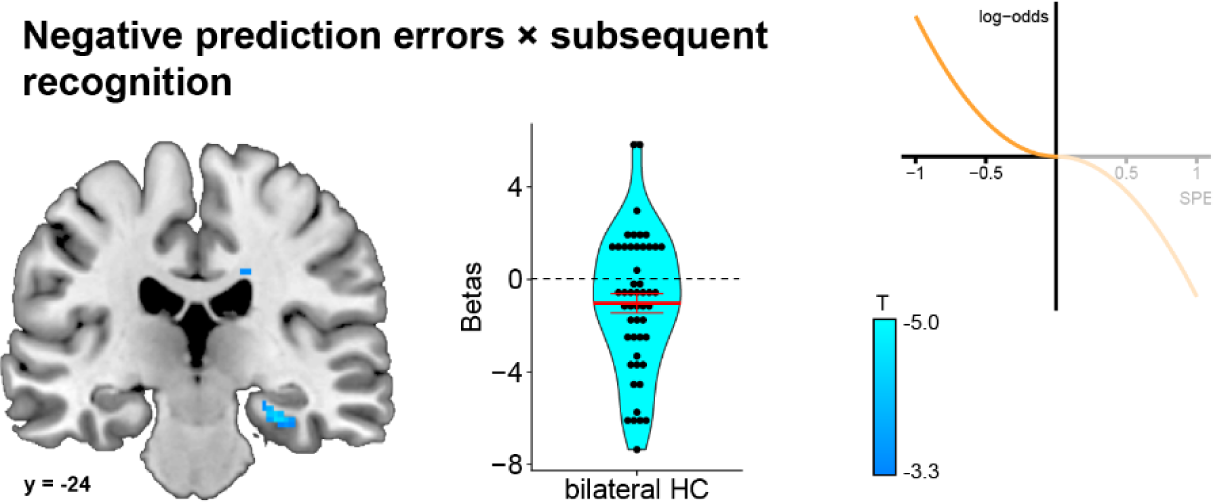
Univariate fMRI analysis of the interaction between negative prediction errors and subsequent memory in the bilateral hippocampus

Particularly for items associated with larger negative PEs that were later recognized, we found decreased BOLD responses in the bilateral hippocampus. Displayed voxels were thresholded at p<.001 for display purposes only. Black dots indicate beta estimates from individual participants, while the red line shows the mean beta estimate over all participants.

### Activity patterns predictive for both PEs and item recognition

Our above analyses showed increased activity for negative PEs in regions of the salience network (dACC and bilateral anterior insula). Regions of the schema network (mPFC, angular gyrus, and precuneus) and the hippocampus, however, showed decreases in activation with larger negative PEs, which was directly linked to improved subsequence recognition. To further elucidate the mechanism through which (negative) PEs facilitate memory formation, we used multivoxel pattern analysis based on activity patterns of areas identified in the univariate analysis to investigate whether a single region would contain both (1) pattern information that can be used to decode PEs and (2) pattern information that predicts subsequent recognition memory (see Methods).

Our results showed that regions associated with negative PEs in the univariate analysis also contained pattern information that enabled us to decode the magnitude of negative PEs (see supplemental Figure S3). Although PEs were decoded above chance already before the outcome presentation which may be due to uncertainty effects, PEs could be decoded best around the time window when the outcome of a trial was revealed and shortly thereafter. The subsequent recognition of an item could only be decoded significantly above chance level with patterns of activity from the insula that occurred around the time the outcome (and by implication, the PE) of trial was revealed (see supplemental Figure S3). However, this effect did not survive a correction for multiple comparisons and therefore needs to be interpreted with great caution.

### PEs are associated with altered connectivity within and between memory-relevant neural networks

Based on the theoretical distinction between ‘standard’ memory processing of events that are in line with prior knowledge and an alternative mode of memory formation for events that are linked to unexpected outcomes, we further hypothesized that items associated with high negative PEs are particularly well remembered because they alter contributions of three main memory networks: (1) the salience network (represented by anterior insula and dACC; Ham et al., 2013; Menon, 2011; Metereau & Dreher, 2013; Seeley et al., 2007), (2) the medial-temporal encoding network (represented by bilateral hippocampus and bilateral parahippocampus), and (3) the schema network (represented by mPFC, precuneus, and angular gyrus; van Kesteren et al., 2012; Vogel et al., 2018a). To address this hypothesis, we analyzed functional connectivity within and between these networks depending on PE magnitudes. For this analysis, we defined a separate GLM with 8 regressors based on combinations of the following factors: onset type (stimulus vs. outcome), outcome (shocked vs. unshocked), and PE magnitude (low if |sPE| < 0.5; high otherwise). After pre-processing the raw times series (see Methods), we based our analysis on the implemented network atlas consisting of several ROIs each to compute within- and between-network correlations (Figure 6A). Here, we focused on the contrast between high and low PEs at the time when the outcome of each trial was revealed. Results showed significant PE-related changes in the connectivity between large-scale networks. Specifically, for large vs. small negative PEs we obtained significantly increased functional connectivity between the salience network and both the schema network (t(49)=2.68, p_corr_=.030 (Bonferroni-corrected), d_av_=0.344) and the medial-temporal encoding network (t(49)=2.18, p=.034, p_corr_=.10 (Bonferroni-corrected), d_av_=0.355; Figure 6B); the connectivity between the schema network and the medial-temporal network did not depend on PEs in unshocked trials, t(49)=0.29, p=.773, p_corr_=1 (Bonferroni-corrected), d_av_=0.046). When we correlated the two PE-related increases in between network connectivity with memory, we found that the increase in functional connectivity between the salience and schema networks was relevant for long-term memory formation, as indicated by its significant correlation with improved hit rates for high negative PE items, r=0.320, t(48)=2.34, p_corr_=.048 (Bonferroni-corrected; Figure 6C); salience-MTEN correlation with hit rates for high negative PE items: r=0.147, t(48)=1.03, p=.31, p_corr_=.62 (Bonferroni-corrected). Furthermore, within-network connectivity decreased for large compared with small negative PEs in the medial-temporal encoding network (t(49)=2.44, p=.018, p_corr_=.055 (Bonferroni-corrected), d_av_=0.307), but not in the salience network (t(49)=1.60, p=.115, p_corr_=.346 (Bonferroni-corrected) d_av_=0.218), nor in the schema network (t(49)=1.24, p=.221, p_corr_=.664 (Bonferroni-corrected), d_av_=0.221; Figure 6D).

**Figure 6.**
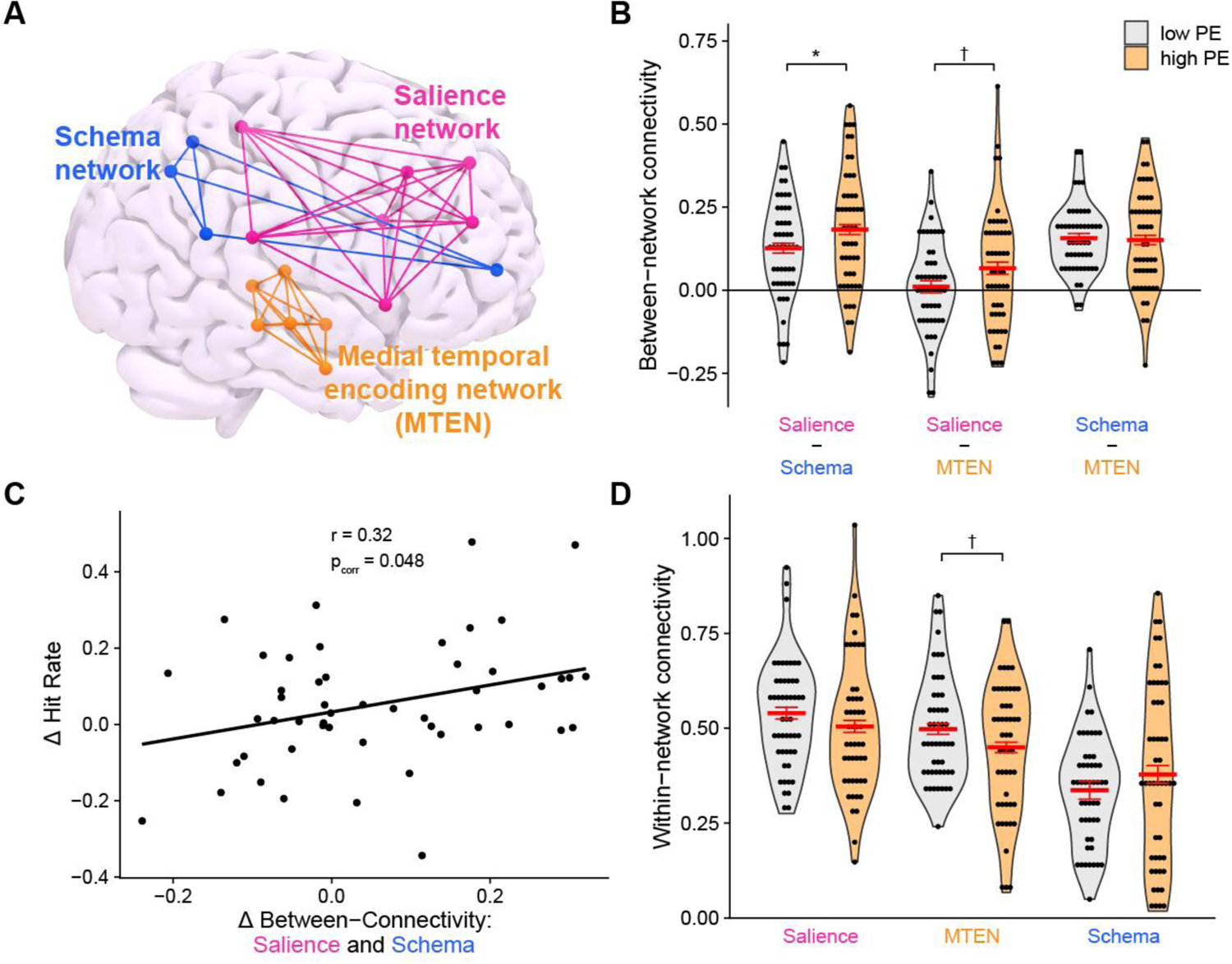
Negative prediction error magnitude is associated with altered within- and between-network connectivity in memory-relevant networks (A) We investigated PE-associated changes in the activity within and between the salience network (rostral prefrontal cortex, supramaginal gyrus, anterior insula, and dACC), schema network (mPFC, precuneus, and angular gyrus) and medial-temporal encoding network (hippocampus and anterior/posterior parahippocampal gyrus). (B) Increases in functional connectivity between salience network and schema network in response to large negative PEs correlated with greater memory enhancement for large negative PEs. (C) Large (vs. small) PEs were associated with significantly increased cross-network connectivity of the salience network with both the schema-network and the medial-temporal encoding network. Thick red bar represent group means, while thin red bars show ± 1 standard error of the mean. (D) Large (vs. small) PEs were associated with significantly decreased within-network functional connectivity in the medial temporal encoding network. † p<.05, * p_corr_<.05 (Bonferroni-corrected).

## DISCUSSION

For decades, PEs have been known to act as teaching signals in reinforcement learning (Cohen, 2008; Schultz, 1998; Sutton & Barto, 1981). However, it was only rather recently discovered that PEs may shape memory formation for episodes preceding the PE event. Here, we combined fMRI with behavioral modelling and large-scale network connectivity analyses to elucidate the mechanisms through which PEs modulate the formation of long-term memories. Our results provide evidence that negative PEs for aversive events promote memory formation for preceding stimuli through a mechanism that is critically distinct from common mechanisms of long-term memory formation and it is tempting to speculate that this mechanism might actively set high PE events apart from events that are in line with prior knowledge and expectation. Critically, the proposed PE-related memory storage mechanism could not be attributed to well-known effects of physiological arousal on memory formation or the effect of a specific prediction itself.

Traditionally, enhanced episodic memory formation has been linked to the medial temporal lobe, including the hippocampus (Davachi & Wagner, 2002; Eichenbaum, 2004; Fernández et al., 1999; Mayes et al., 2007; Reed & Squire, 1997; Shrager et al., 2008). The negative PE-related memory enhancement, however, was not linked to enhanced but even to decreased hippocampal activity. Further, when participants experienced a negative PE, there was a reduction in connectivity within the medial-temporal encoding network. While hippocampal activity was reduced for negative PEs, we obtained significantly increased activity in the anterior insula and dACC for negative PE events. Both of these regions have previously been implicated in error monitoring, conscious perception of errors, and aversive PE signaling (Bastin et al., 2016; Fazeli & Büchel, 2018; Garrison et al., 2013; Preuschoff et al., 2008; Taylor et al., 2007; Ullsperger et al., 2010). Moreover, both the anterior insula and the dACC are key regions of the salience network (Ham et al., 2013; Menon, 2011), which signals biologically relevant events and the need for a behavioral or cognitive change (Dosenbach et al., 2006; Kerns, 2004). Furthermore, the salience network has been proposed to dynamically change the control of other large-scale networks (Sridharan et al., 2008). In line with this idea, we obtained here a trend for increased functional connectivity between the salience network and the medial-temporal encoding network for negative PEs.

In addition to the negative PE-related decrease in hippocampal activity, there was also a marked decrease in the activity of angular gyrus, precuneus, and mPFC for events associated with negative PEs. Together, these areas form a ‘schema-network’, in which the mPFC is thought to detect a congruency of events with prior knowledge and to then integrate these events into existing knowledge representations (van Kesteren et al., 2012; Vogel et al., 2018b). This schema-network is of particular interest because PEs signal that an event differs from prior experience. When the organism experiences large PEs, this indicates that new information conflicts with prior knowledge and should therefore be stored separately from existing schema-congruent memories (van Kesteren et al., 2012). This idea is supported by the obtained negative PE-associated decrease in areas constituting the schema network.

Moreover, there was also increased connectivity between the salience network and the schema-network when individuals experienced a negative PE and this PE-related change in large-scale network connectivity was directly correlated with the negative PE-driven memory enhancement. Together these findings suggest that the negative PE-induced enhancement of episodic memory is not driven by an enhancement of common medial temporal mechanisms of memory formation but by a distinct mechanism that is linked to the salience network and separates PE events from experiences that are in line with prior knowledge.

The salience network has often been related to physiological arousal (Xia et al., 2017; Young et al., 2017) which is well known to mediate the superior memory for emotionally arousing events (Cahill & McGaugh, 1998; McGaugh, 2018). Although one might assume that high negative PEs may have elicited arousal which then enhanced memory storage, our data speak against this alternative and suggest that negative PE-related memory enhancement was not due to increased physiological arousal. First, aversive shocks per se had no influence on memory formation. Moreover, in our trial-level behavioral models, neither anticipatory nor outcome-related SCRs as measures of physiological arousal predicted memory formation.

Only in a combined model featuring additionally uncertainty and PEs, larger outcome-related SCRs were linked to improved recognition performance. Even in this combined model, we still found clear evidence for complementary effects of PEs (and uncertainty) beyond arousal measures. Importantly, specific neural clusters associated with negative PEs were identified in a model that controlled for physiological arousal. These results indicate that the effects of PEs on episodic memory formation cannot be explained by traditional arousal-based models.

Further, although greater shock expectancy was by itself linked to enhanced memory formation, we found that effects of PEs, which contrast such expectations with observed outcomes, explained recognition beyond main effects of shock expectations and observed outcomes. This speaks against an alternative account of our findings in which the mere prediction of an aversive event, possibly through increased attention to the predictive stimulus, is sufficient to explain our observed effects on memory formation. Instead, PEs, which indicate that the current model of the world might need to be revised, seem to contribute to memory formation beyond isolated effects of expectation or outcomes.

It is also important to note that our findings go above and beyond previous results showing an enhanced memory for novel or surprising stimuli (Cycowicz & Friedman, 2007; Strange & Dolan, 2004). We show here that, rather than the novelty of a stimulus, the discrepancy between expected and experienced consequences of a stimulus affected its memorability. This is particularly remarkable as these consequences were only revealed after a stimulus had already disappeared, thus ruling out a simple increase of attentional processing.

While our findings show that PEs may promote memory formation, this effect depended on the direction of the PE. Previous behavioral findings could not differentiate effects of negative and positive PEs in an aversive context (Kalbe & Schwabe, 2020) and studies on the role of reward-related PEs yielded inconsistent findings as to whether the direction of the PE matters for episodic memory formation (Ergo et al., 2020; Jang et al., 2019; Rouhani et al., 2018). By including CS categories that were linked to either a high or a low shock probability, we obtained relatively symmetrical distributions of positive and negative PEs. This allowed us to investigate the distinct effects of positive and negative PEs. Interestingly, we found that memory effects depended on the sign of PEs, with negative PEs being associated with better recognition performance and larger positive PEs showing opposite, negative effects on recognition performance. In other words, not only PE magnitude affected memory formation, but also their direction, that is, whether shock probabilities were under- or overestimated. The neural signature of positive PEs was clearly distinct from the neural underpinnings of negative PEs. Positive PEs were associated with clusters of increased activation in the left superior parietal lobule and the right middle temporal gyrus and decreased activation of the left supramarginal gyrus. The superior parietal lobe has been linked to internal representations of sensory inputs before (Wolpert et al., 1998) as well as to contralateral sensorimotor coding of body parts (Wolbers, 2003). As the electric shock was applied to the right leg and increased superior marginal activation was observed in the left hemisphere, the observed activity pattern might point to increased processing of the electric shock. Furthermore, the supramarginal gyrus has been previously associated with motor planning (Potok et al., 2019) and unexpected somatosensory feedback perturbation (Golfinopoulos et al., 2011). Thus, it is tempting to speculate that positive PEs resulted in more pronounced processing of the (unexpected) electric shock, which distracted from the mnemonic processing of the encoded stimulus and hence led to decreased subsequent recognition memory.

Closely related but conceptually distinct from PEs is prediction uncertainty. While PEs only become apparent after an outcome has been revealed, uncertainty emerges as soon as a potentially threatening stimulus is presented. Moreover, whereas PEs are calculated by contrasting predictions with following outcomes, uncertainty is independent of outcomes. Our design allowed us to distinguish between PEs and uncertainty. Same as for positive PEs, we found that uncertainty about the possible occurrence of a shock was associated with decreased recognition performance. At the neural level, uncertainty was paralleled by decreased activation in bilateral medial occipital areas, possibly reflecting diminished visual processing of stimuli associated with uncertain outcomes, which might explain the uncertainty-related impairment in recognition. In addition, uncertainty was associated with reduced mPFC activation, a region implicated in beliefs and the inference of hidden states (Starkweather et al., 2018; Yoshida & Ishii, 2006).

In summary, we provide behavioral and neural evidence for a critical impact of PEs on long-term memory formation for events preceding the PE, thereby bridging the traditionally separated fields of associative learning and long-term memory. In addition to the magnitude of the PE, our results show that the direction of the PE affects memory formation. Whereas positive PEs reduced subsequent memory, negative PEs promoted memory formation. In particular for negative PEs, our results suggest a qualitative shift in the contributions of large-scale neural networks to memory formation. Negative PEs reduced the processing of events in the schema network and the medial-temporal encoding network both of which are involved in ‘standard’ long-term memory formation. Instead, such schema-incongruent experiences might be particularly well remembered because they are encoded distinctly from more mundane experiences, perhaps at an exemplar-level, in a process that is likely mediated through the salience network. Importantly, these memory enhancements and related neural changes could not be explained by the prediction itself or mere changes in physiological arousal, thus pointing to a rather ‘cognitive’ mechanism of memory enhancement. These findings may have relevant implications for the treatment of fear-related mental disorders, suggesting that it might be beneficial to explicitly activate patients’ negative outcome expectations prior to the exposure to the feared stimulus, as the absence of the feared consequence in the therapeutic context should produce strong fear-incongruent memories. More generally, our results provide novel insights into the mechanisms underlying the exceptional memory for episodes in the context of unexpected events, such as meeting Barack Obama in the supermarket.

## METHODS

### Participants

Sixty-one healthy volunteers (35 women, 26 men; mean age±SD=24.97±4.65 years) participated in this experiment. Eleven participants had to be excluded from analysis due to excessive head motion in the scanner (>5mm within a single experimental block; N= 2), incidental finding of a frontal lesion (N=1), missing >25% of responses on the task (N=6), selecting only extreme ratings (i.e., 0% and 100%; N=1), or not returning for the second experimental day (N=1). To determine the target sample size, we performed an a-priori power analysis based on previous findings of binary aversive PE effects on episodic memory formation (Kalbe & Schwabe, 2020). As this study used a conceptually similar generalized linear mixed-effect model, we applied a simulation-based approach using the *SIMR R* package (Green & MacLeod, 2016). We assumed the same effect size but increased the number of trials from 60 to 120 to account for the modified design in the present study. This indicated that a sample size of N=50 participants would result in a statistical power of above .95. All participants met safety criteria for MRI and electrodermal stimulation, had normal or corrected-to-normal vision, were right-handed, had never studied psychology nor neuroscience, did not suffer from any psychiatric or neurological conditions, and reported no alcohol abuse, nor use of any illicit drug. They were paid 45€ upon completion of the second experimental day. The study protocol was approved by the ethics board of the University of Hamburg and all participants provided informed consent prior to their participation.

### Experimental procedure

The experiment took place on two consecutive days. On the first experimental day, participants completed a combined incidental encoding and fear learning task in the MRI. About 24 hours later, they completed a surprise recognition test for stimuli presented during the encoding session. At the beginning of the first experimental day, participants provided informed consent and were prepared for the MRI scanner by placing a pair of MRI-safe gelled disposable electrodes (BIOPAC systems, Goleta, CA, USA) over the thenar eminence of the left hand to measure skin conductance as an indicator of physiological arousal during the encoding task using the BIOPAC MP-160 system (BIOPAC systems, Goleta, CA, USA). Another pair of electrodes was placed on the right side of the right lower leg, approximately 20cm above the ankle, and used to administer aversive electric shocks during the fear learning task. Shocks were applied using the BIOPAC STMISOC (BIOPAC systems, Goleta, CA, USA) connected to a BIOPAC STM100C stimulator (BIOPAC systems, Goleta, CA, USA). After participants were placed in the scanner, they first completed an unrelated task that included stimuli that were critically distinct from the stimuli used in this experiment.

Prior to the start of the fear learning task, shock intensity was adjusted to be unpleasant but not painful by administering a series of test shocks that increased in intensity until participants rated the shocks as not yet painful but highly unpleasant. Participants then received detailed written instructions about the following fear learning task. On each trial, participants saw an image that was presented centrally on a screen for 4.5s (Figure 1A).

Beneath each image, participants saw a slider that always started at 50% and could be adjusted to any integer value between 0% and 100% by using the left and right buttons of an MRI-compatible response box (Current Designs Inc., Philadelphia, USA). Participants were instructed that while each image was present, they should adjust the slider to a value that corresponded with their prediction of the probability that a shock would follow. Participants were requested to confirm their rating by pressing the central button on the response box. In 40 out of the total of 120 trials, a 200ms shock to the right lower leg followed immediately after image offset. Between trials, there was a jittered white fixation cross presented for 5s to 8s. This relatively long inter-trial interval allowed us to observe the slowly emerging SCR in response to each outcome as well as to separate trials at the neural level. Critically, the probabilities of a shock were linked to image categories. While participants were explicitly instructed that they would see images of vehicles, clothing, and tools, they were not told that these categories would be linked to pre-defined shock contingencies. Participants were informed that their predictions would have no effect on the probability that a shock would occur, but that their aim should still be to improve their predictions over the course of the task. Out of 40 occurrences of the CS^a+^ category, 27 were followed by a shock, corresponding to a shock probability of approximately 2/3. Likewise, 40 occurrences of the CS^b+^ category were followed by a shock in 13 trials, leading to a shock probability of approximately 1/3 for the CS^b+^. Finally, the 40 occurrences of the CS-category were never followed by a shock.

The six possible combinations of image categories (i.e., vehicles, clothing, tools) with conditioning categories (i.e., CS^a+^, CS^b+^, CS^-^) were counterbalanced across participants. Participants completed four blocks with 30 trials each, resulting in a total of 120 trials. Between blocks, participants had the opportunity to ask the experimenter to slightly reduce the shock intensity in cases when shocks had become painful.

After an interval of 22h to 26h, participants returned for a surprise recognition test outside of the MRI scanner. In this recognition test, they saw all 120 images that had been presented on the previous day randomly intermixed with the same number of previously unseen (‘new’) images from the same three categories (40 new images per category). For each image, participants had a maximum of 6s to indicate whether the current image had been presented on the previous day (‘old’) or not (‘new’) and how confident they were, using buttons corresponding to ‘definitely old’, ‘maybe old’, ‘maybe new’, and ‘definitely new’. Between each of the 240 trials of the recognition test (120 old, 120 new), a white fixation cross appeared centrally for 1 to 2s.

### MRI data acquisition

Functional MRI data were acquired during the incidental encoding session on a Siemens Magnetom Prisma 3T scanner equipped with a 64-channel head coil. For each of the four functional runs, approximately 185 volumes were recorded using a multi-band echo-planar imaging (EPI) sequence with the following parameters: 60 axial slices of 2mm depth, slice orientation parallel to the AC-PC line, phase-encoding in AP direction, repetition time (TR) of 2000ms, echo time (TE) of 30ms, 60-degree flip angle, 224mm × 224mm field of view (FOV), 2mm isotropic resolution, EPI factor of 112, echo distance of 0.58ms. For each block, four images were recorded before the start of the behavioral task to ensure equilibrium magnetization. These initial images were discarded as dummy scans during further analyses. Following the last functional run, a T1-weigthed scan was acquired with 256 slices, coronal orientation, repetition time (TR) of 2300ms, echo time (TE) of 2.12ms, a 240mm x 240mm field of view (FOV), and a 0.8mm × 0.8mm × 0.9mm voxel size.

### Behavioral analysis

For each individual trial, the prediction uncertainty (*PU*) was derived from participants’ shock predictions, while signed PEs (*sPE*) were calculated by contrasting predictions with actual outcomes. Specifically, the *PU* is a continuous variable that can take any value between 0 (least possible uncertainty) and 1 (maximum uncertainty) and was calculated as:

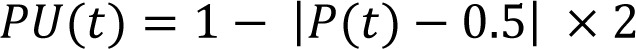

Where *P(t)* is the continuous explicit shock prediction made by the participant in trial *t* (ranging from 0 to 1).

The *sPE* in trial *t* is a continuous variable that can take any value between −1 and +1 and was calculated as:

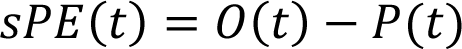

Where *O(t)* is the binary outcome in trial *t* (coded 0 when no shock occurred and 1 when a shock occurred). Note that the sign of the *sPE* contains information about the outcome of the trial. *sPEs* <0 could only occur in unshocked trials, while *sPEs* >0 could occur when a shock occured. Only for *sPE*=0, the binary outcome of the trial is ambiguous.

The prediction uncertainty *PU(t)* for any trial *t* can also be calculated directly from the *sPE* (but not vice versa) using:

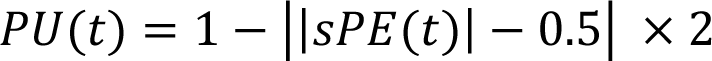

To test influences of uncertainty, PEs, and arousal (measured through SCRs) on episodic memory formation, we performed mixed-effects logistic regression at the level of individual trials, as implemented in the *lme4 R* package (Bates et al., 2015). The binary recognition of a previously presented item (collapsed over confidence ratings) was treated as the dependent variable, coded 0 for misses and coded 1 for hits. Following recommendations to maximize the generalizability of these models (Barr et al., 2013), we included the maximum random effects structure, estimating random intercepts and random slopes per predictor per subject. We did not include random intercepts per item to account for different baseline memorability as their inclusion led to singular fit in some models.

### Skin conductance analysis

During the incidental encoding session of the first experimental day, we recorded electrodermal activity as a measure of physiological arousal. These data were analyzed in Ledalab Version 3.4.9 (Benedek & Kaernbach, 2010) using a Continuous Decomposition Analysis (CDA) to derive the average phasic driver within given response windows. In short, the CDA aims to separate the continuous skin conductance data into a tonic, stimulus-independent component, and a phasic, stimulus-driven component. To obtain more precise estimates of the underlying sudomotor nerve activity compared with more traditional methods such as a through-to-peak analysis, the CDA only considers changes in the phasic component in response to an event. As a first measure, we defined anticipatory SCRs as reactions occurring from the onset of the decision in each trial (i.e., the confirmation of the shock rating) until the end of the stimulus presentation (i.e., exactly 4.5s after stimulus onset).

Additionally, we defined outcome-related SCRs to occur 0.5s after the outcome of the current trial was revealed (i.e., whether a shock would occur or not) until 2.9s after the outcome onset to ensure that this measure would capture activity evoked by the current trial, but not the following. Skin conductance data were downsampled from 1000Hz to 50Hz and optimized using four sets of initial values. The minimum amplitude threshold was set to 0.01µS for both anticipatory and the outcome-related SCRs. Individual physiological factors, such as the thickness of the corneum can greatly affect the range of observed SCRs (Figner & Murphy, 2011). To account for this interindividual variability, both the anticipatory and the outcome-related SCRs were standardized by dividing the average phasic driver estimate by the maximum average phasic driver value observed in any trial.

### fMRI preprocessing

Functional MRI data were preprocessed in MATLAB using SPM12 (https://www.fil.ion.ucl.ac.uk/spm/software/spm12). First, functional volumes were spatially realigned to the first image in the time series. This step also yielded six motion parameters used in univariate analyses to control for motion-related activation artifacts. Realigned volumes were co-registered to each participant’s structural image. Then, images were spatially normalized into standard stereotactic (MNI) space using unified segmentation. For univariate fMRI analyses, the normalized functional images were additionally smoothed with an 8mm full-width half-maximum Gaussian kernel. For multivoxel pattern analysis, the unsmoothed normalized images were used.

### Univariate fMRI analyses

Based on results from behavioral modelling, we identified (1) quadratic prediction uncertainty, (2) quadratic signed PEs, and (3) physiological arousal (measured through anticipatory and outcome-related SCR, respectively) as key variables to explain episodic memory formation in this fear learning task. To investigate the neural basis of these effects, we modelled the fMRI time series using generalized linear models (GLMs). These models included regressors of the onsets of the stimulus and outcome presentation as predictors of interest, and nuisance regressors to account for head movement (i.e., the six movement parameters derived from spatial realignment. As behavioral data suggested separable effects of positive vs. negative quadratic prediction errors on memory formation, we fitted separate models for unshocked trials (corresponding to negative PEs) and shocked trials (corresponding to positive PEs). Both models featured onsets of stimuli with shock expectancy and prediction uncertainty as parametric modulators. To control for possible effects of arousal, standardized anticipatory SCRs were also placed as an additional parametric modulator on stimulus onsets. A second regressor featured onsets of with quadratic prediction errors as the critical parametric modulator. Again, we controlled for possible confounding effects of arousal by placing standardized outcome-related SCRs as an additional parametric modulator on outcome onsets.

For the estimation procedure, data from each of the four experimental blocks were concatenated using the spm_fmri_concatenate function in SPM12, a high-pass filter at 1/128Hz was applied, and an AR(1) process was used to adjust for temporal autocorrelation. Second-level analysis were constructed from each subject’s first level contrasts using a standard one-sample t-test approach in SPM. We thresholded all resulting t-maps using a whole-brain voxel-level family-wise error corrected P value of PFWE<.05.

To link the memory-promoting effects of negative PEs with subsequent recognition, we specified an additional univariate fMRI model with onsets of unshocked outcomes as a regressor and PEs (ranging between 0 and 1), the binary subsequent recognition of an item (coded 0 for misses and 1 for hits) and their interaction as parametric modulators. This model was estimated using the same procedure as described above. Based on results from the univariate fMRI analysis of negative PEs, we defined the bilateral insula, the ventromedial prefrontal cortex, and the bilateral hippocampus as three regions of interests. Voxels belonging to each of these regions were identified based on an existing anatomical atlas (Harvard-Oxford structural atlas; Desikan et al., 2006). Within each region of interest, we obtained the average estimated beta weight per participant and performed one sample t-tests to test whether these were significantly different from 0. Our analysis focused on the interaction between PEs and subsequent recognition, as this specific interaction links the processing of PEs with its effect in memory formation.

### Multivoxel pattern analyses (MVPA)

To test which of the regions identified in the univariate analysis contained pattern information about (1) the extent of PEs and (2) the probability that an encoded item would later be recognized, we performed a multivoxel pattern analysis (MVPA; Kriegeskorte, 2011). As PEs are continuous, decoding them from neural data constitutes a regression problem, while the later recognition of an item is binary, and its decoding therefore constitutes a classification problem. Hence, these problems required slightly different machine learning algorithms, although both were selected from the class of support vector machines and the general data preparation and model fitting procedure was very similar for both problems.

The MVPA was performed on t-maps that were generated in SPM12 from the unsmoothed, normalized functional data from each participant. Separate generalized linear models were estimated to extract several trial-specific t-maps of each of the following points in time relative to each outcome: −4, −2, 0, 2, 4. Extracting multiple activation maps per trial in this way allowed us to address specifically the question when exactly relevant pattern information was present. Note that stimulus onsets were always 4.5s before outcome onsets and were therefore represented by the offset −4. Also note that the temporal distance of 2s between offsets corresponds with the TR of the EPI sequence. In each GLM, each single onset of the outcome event, offset by the currently estimated point in time, was entered as its own regressor. Therefore, for each trial and offset, we generated unique beta-maps, which were then transformed to t-maps to normalize them (Misaki et al., 2010). The GLM used the same parameters as in the univariate analyses, namely, concatenation of experimental blocks, a high-pass filter at 1/128Hz, and an AR(1) process to adjust for temporal autocorrelation.

T-maps representing individual trials and offsets per participant were then further processed in Python 3 using the Nilearn module (Abraham et al., 2014). Whole-brain t-maps were masked with ROIs identified in the univariate analysis. Specifically, in the case of larger regions (e.g., insula), we created new masks by identifying the peak voxel per region from the second level analysis of univariate results reported earlier and including voxels within a 6mm-radius of each peak voxel. For the bilateral hippocampus, we used existing anatomical masks (Harvard-Oxford structural atlas; Desikan et al., 2006).

To prepare extracted data from each ROI for use with common machine learning algorithms, the 4-dimensional t-maps (three spatial and one temporal dimension) were reshaped to a samples-by-features matrix (number of trials × number of voxels in ROI). Further, data were z-standardized using the StandardScaler implementation in scikit-learn (Pedregosa et al., 2011). To predict PEs from neural data, we trained a support vector regression (SVR) with a linear kernel as implemented in scikit-learn with the regularization hyperparameter *C* fixed at 1 and the negative mean squared error as the performance metric. Similarly, to predict the binary recognition of an item, we trained a linear support vector classifier using the LinearSVC implementation in scikit-learn with the regularization parameter *C* fixed at 1 and the area under the receiver operating characteristic (ROC) curve as the performance metric to account for imbalanced classes due to uneven numbers of hits and misses for each participant. For both decoding tasks, we used leave-one-block-out cross-validation to evaluate decoding performance, such that three blocks were always used for training and the remaining block was used for validation. Performance metrics from all four possible training-validation combinations were averaged to compute the mean performance.

To establish a baseline performance at chance level that can be used to compare each fitted model against, for each “true” performance score, we also performed the exact same preprocessing and training procedure using 100 separate random permutations of the true labels as a permutation test (Nichols & Holmes, 2002). Therefore, the above-chance-performance of a predictive model could be conceptualized here as the distance between the performance achieved with true labels and the mean performance in the permutation test.

### Network-connectivity analyses

We performed analyses of functional connectivity in the CONN toolbox (Whitfield-Gabrieli & Nieto-Castanon, 2012) to assess how within- and between-network connectivities of memory-relevant brain networks differed depending on PE magnitudes. As this analysis did not allow for continuous parametric modulators, we instead split PEs into low (|sPE|<0.5) vs. high (|sPE|≥0.5). Our analyses focused on PE effects at outcome time for unshocked trials.

However, in the specific GLM for this analysis, we included onset regressors for each combination of the following factors: stimulus vs. outcome onsets, shocked vs. unshocked, and low vs. high PEs. This resulted in a total of 8 regressors in this model. In a first-level analysis, to denoise data, we applied a linear detrending and a standard band-pass filter of 0.008 to 0.09 Hz. Besides the just mentioned effects of PEs, we added white matter, cerebrospinal fluid, and movement regressors obtained from spatial realignment as additional confounds to the model. Further analysis focused on pre-defined regions of interest and networks implemented in the CONN toolbox: (i) dorsal anterior cingulate cortex, bilateral anterior insula, bilateral rostral prefrontal cortex and bilateral supramaginal gyrus forming the *salience network* (Menon, 2011); (ii) medial prefrontal cortex, bilateral angular gyrus and precuneus forming the schema network (van Kesteren et al., 2012; Vogel et al., 2018a); and (iii) bilateral hippocampus, bilateral anterior parahippocampal gyrus, and bilateral posterior parahippocampal gyrus as the medial temporal encoding network (Fernández et al., 1999; Shrager et al., 2008).

## Acknowledgements

We gratefully acknowledge the support of Friederike Baier, Jan-Ole Großmann, Vincent Kühn, and Ricarda Vielhauer during data collection.

## Author contributions

Conceptualization, F.K. and L.S.; Methodology, F.K. and L.S.; Formal analysis, F.K.; Investigation, F.K.; Writing –Original Draft, F.K. and L.S.; Writing –Review & Editing, F.K. and L.S.; Funding Acquisition, L.S.; Resources, L.S.; Supervision, L.S.

## Data availability

Behavioral, SCR, and fMRI data that support the findings of this study are available at OSF: https://osf.io/3atyr/.

## Code availability

Custom code used to analyze and model the data is available at OSF: https://osf.io/3atyr/.

## SUPPLEMENTAL MATERIAL

**Supplemental Figure 1.**
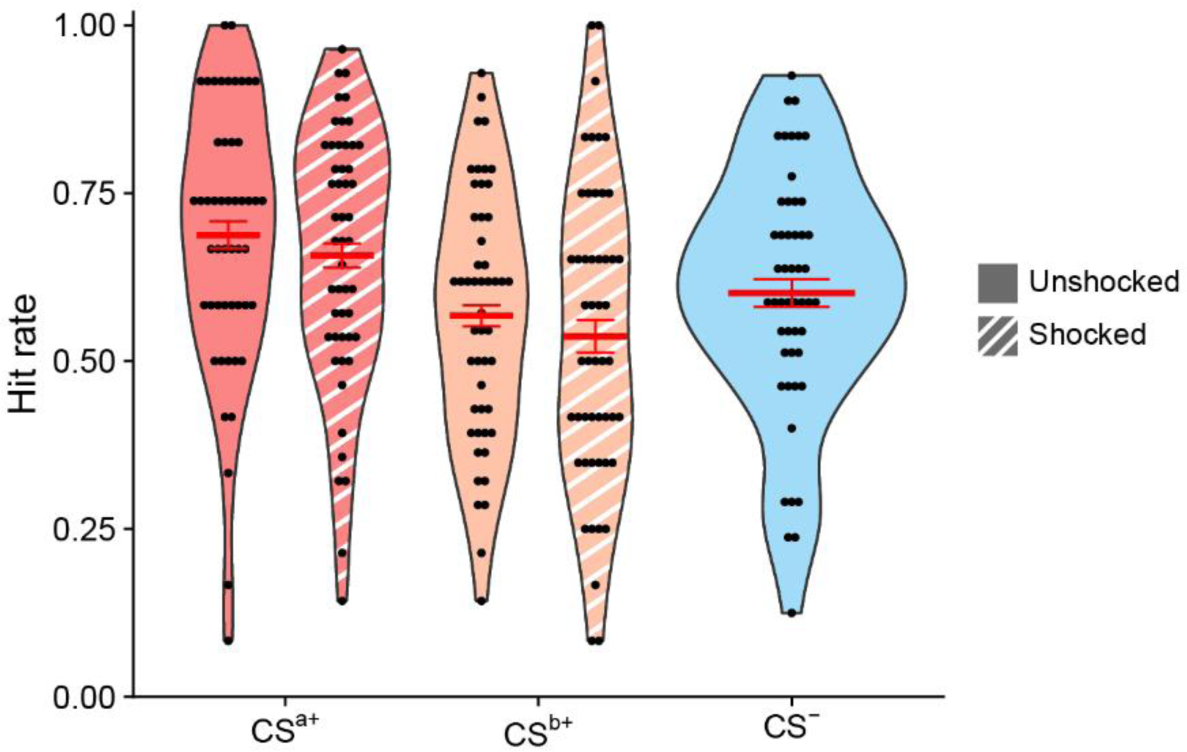
Hit rates by CS category and trial outcome. Hit rates for items from the CS^a+^ category were larger compared with both CS^b+^ and CS-items. Importantly, the greater number of aversive shocks for CS^a+^ items could not explain this memory difference, as participants descriptively even showed slightly decreased hit rates for shocked (compared with unshocked) items in both the CS^a+^ and CS^b+^ category.

**Supplemental Figure 2.**
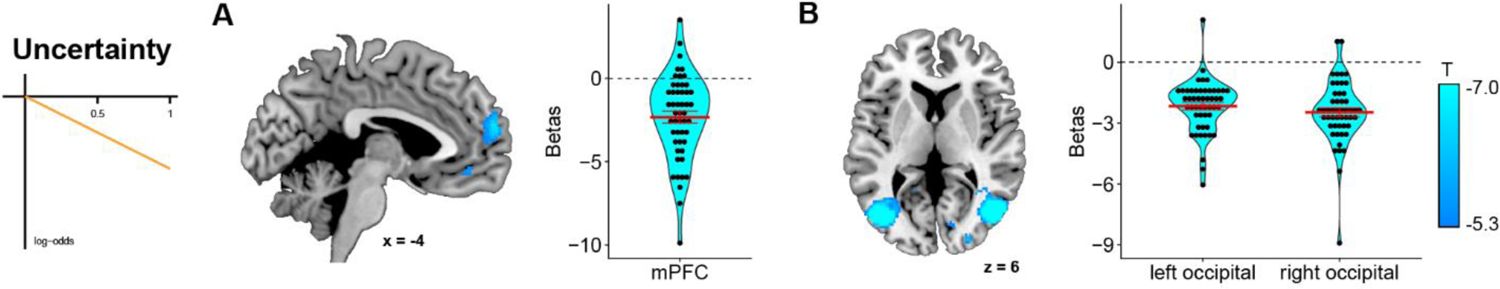
Univariate fMRI analysis to identify regions associated with uncertainty in unshocked trials For prediction uncertainty in unshocked trials, we obtained decreased BOLD responses in the mPFC and bilateral occipital areas (A, B). Only voxels significant at 5%-level after whole-brain family-wise error (FWE) correction (peak level) are displayed. Black dots indicate beta estimates from individual participants, while the red line shows the mean beta estimate over all participants. Thin red bars show ± 1 standard error of the mean.

**Supplemental Figure 3.**
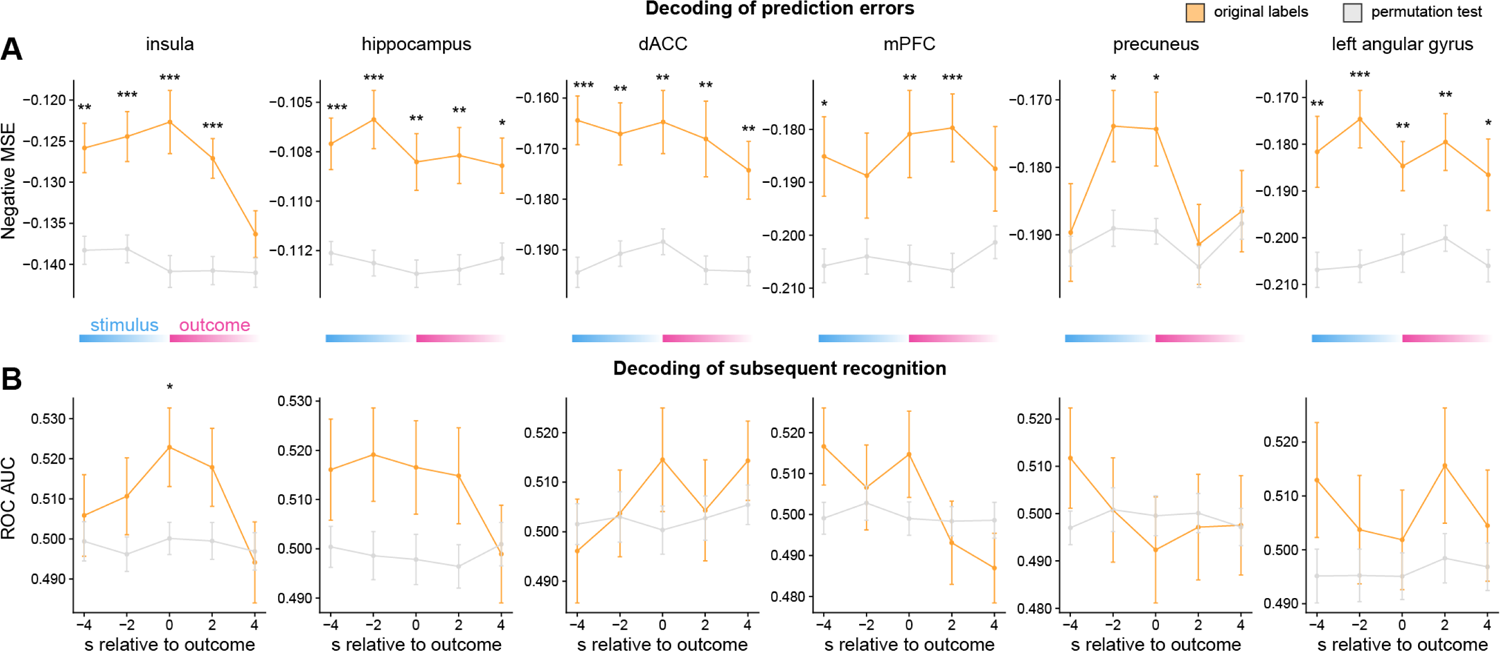
Decoding of prediction errors and subsequent recognition using multivoxel pattern analysis (MVPA) (A) Activity patterns from regions associated with negative PEs in the univariate analysis could be used to decode the magnitude of negative PEs. Best decoding performance was generally achieved around the time the outcome was revealed. (B) Subsequent recognition of an item could be decoded significantly above chance level using patterns of activity from the insula specifically when the outcome of a trial was revealed. * p<.05, ** p<.01, *** p<.001.

**Supplemental Table 1.** Univariate fMRI results for PEs

| Effect | Direction | Region | x | y | z | T | p(FWE) |
| --- | --- | --- | --- | --- | --- | --- | --- |
| Negative PEs<br>(cluster extend threshold: $k \geq 10$) | positive | right anterior insula | 32 | 22 | -8 | 8.51 | $2.09 \times 10^{-6}$ |
| | | left anterior insula | -34 | 18 | 4 | 8.01 | $9.97 \times 10^{-6}$ |
| | | dACC | 8 | 26 | 42 | 7.63 | $3.39 \times 10^{-5}$ |
| | | right inferior frontal gyrus | 48 | 26 | 22 | 5.68 | $1.43 \times 10^{-2}$ |
| | | right middle frontal gyrus | 42 | 12 | 30 | 5.52 | $2.31 \times 10^{-2}$ |
| | negative | left hippocampus | -26 | -20 | -16 | 8.49 | $2.20 \times 10^{-6}$ |
| | | precuneus | -12 | -56 | 18 | 8.41 | $2.89 \times 10^{-6}$ |
| | | mPFC | -8 | 50 | -6 | 8.27 | $4.45 \times 10^{-6}$ |
| | | left angular gyrus | -44 | -70 | 26 | 7.86 | $1.64 \times 10^{-6}$ |
| | | right hippocampus | 24 | -22 | -18 | 7.11 | $1.73 \times 10^{-4}$ |
| | | left middle temporal gyrus | -62 | -12 | -16 | 6.61 | $8.29 \times 10^{-4}$ |
| | | left superior frontal gyrus | -24 | 36 | 50 | 6.05 | $4.74 \times 10^{-3}$ |
| | | left primary motor cortex | -28 | -28 | 60 | 5.85 | $8.66 \times 10^{-3}$ |
| | | left caudate | -8 | 24 | 10 | 5.74 | $1.20 \times 10^{-2}$ |
| | | right middle temporal gyrus | 60 | -4 | -22 | 5.58 | $1.94 \times 10^{-2}$ |
| Positive PEs<br>(no cluster extend threshold) | positive | left superior parietal lobule | -32 | -72 | 52 | 6.05 | $3.01 \times 10^{-3}$ |
| | | left lateral temporal cortex | -64 | -40 | -8 | 5.47 | $1.72 \times 10^{-2}$ |
| | negative | left supramarginal gyrus | -60 | -30 | 40 | 5.40 | $2.10 \times 10^{-2}$ |

All displayed peaks were significant at $p < .05$ after whole-brain voxel-level family-wise error correction. Additional minimal cluster extend thresholds of $k \leq 10$ were only applied were indicated.

**Supplemental Table 2.** Univariate fMRI results for uncertainty

| Effect | Direction | Region | x | y | z | T | p(FWE) |
| --- | --- | --- | --- | --- | --- | --- | --- |
| Uncertainty (unshocked trials) | positive | right middle frontal gyrus | 46 | 24 | 34 | 6.08 | $5.27 \times 10^{-3}$ |
| | negative | right extrastriate cortex | 50 | -70 | 6 | 11.89 | $3.76 \times 10^{-11}$ |
| | | left extrastriate cortex | -40 | -74 | 4 | 11.76 | $5.63 \times 10^{-11}$ |
| | | left postcentral | -44 | -30 | 62 | 8.38 | $3.88 \times 10^{-6}$ |
| | | right cuneus | 20 | -82 | 40 | 7.49 | $6.53 \times 10^{-5}$ |
| | | right lingual gyrus | 16 | -62 | -4 | 7.09 | $2.29 \times 10^{-4}$ |
| | | left lingual gyrus | -14 | -72 | 0 | 7.01 | $2.90 \times 10^{-4}$ |
| | | mPFC | -4 | 62 | 18 | 6.76 | $6.46 \times 10^{-4}$ |
| | | left middle temporal gyrus | -64 | -16 | -10 | 6.71 | $7.61 \times 10^{-4}$ |
| | | dorsal posterior cingulate area | -10 | -24 | 46 | 6.67 | $8.67 \times 10^{-4}$ |
| | | right primary visual cortex | 12 | -78 | 2 | 6.56 | $1.18 \times 10^{-3}$ |
| | | left temporal pole | -58 | 4 | -24 | 6.39 | $2.06 \times 10^{-3}$ |
| | | right fusiform area | 42 | -28 | -16 | 6.37 | $2.13 \times 10^{-3}$ |
| | | inferior temporal area | 24 | -72 | 28 | 6.34 | $2.35 \times 10^{-3}$ |
| | | right anterior cingulate area | 8 | -12 | 50 | 6.34 | $2.37 \times 10^{-3}$ |
| | | right superior parietal lobule | 28 | -54 | 62 | 6.21 | $3.55 \times 10^{-3}$ |
| | | left fusiform area | -40 | -46 | -24 | 5.85 | $1.07 \times 10^{-2}$ |

All displayed peaks were significant at $p < .05$ after whole-brain voxel-level family-wise error correction and clusters extended a threshold of $k = 10$ voxels. For shocked trials, there were no significant voxels under this criterion.

**Supplemental Table 3.**
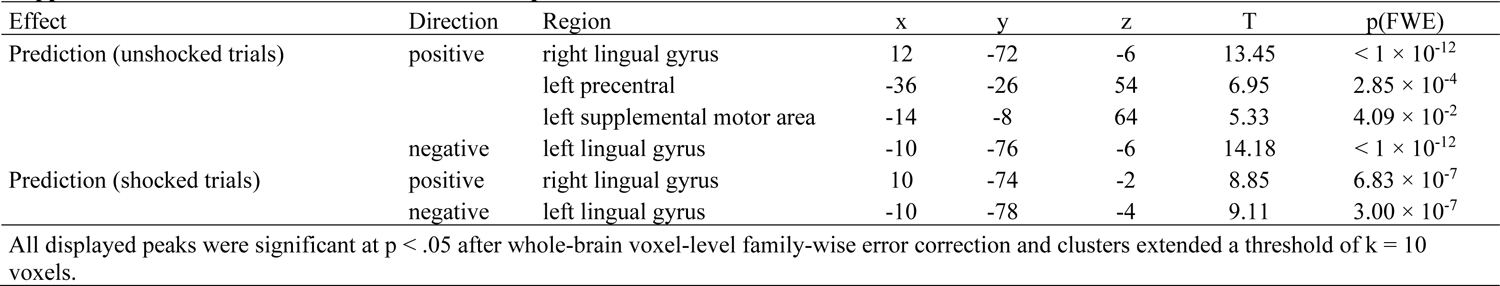
Univariate fMRI results for predictions

